# Repurposing efavirenz, the HIV antiretroviral drug for Chikungunya virus infection

**DOI:** 10.1101/2023.09.29.560149

**Authors:** Sanketkumar Nehul, Ruchi Rani, Prasan Kumar Panda, Pravindra Kumar, Shailly Tomar

## Abstract

Chikungunya virus (CHIKV) has frequently recurred in recent decades, causing outbreaks worldwide in tropical and subtropical regions. The re-emergence of CHIKV poses a substantial risk to human health as no efficacious drugs are currently available to curb new outbreaks. Here, the anti-CHIKV activity of efavirenz was investigated by *in vitro* cell culture-based antiviral assay, immunofluorescence assay (IFA), and quantitative reverse transcription polymerase chain reaction (qRT-PCR). Efavirenz is a non-nucleoside reverse transcriptase inhibitor (NNRTI) used for treatment of acquired immunodeficiency syndrome (AIDS), and it has good oral bioavailability, long half-life and affordable low cost. This study demonstrated dose-dependent robust anti-CHIKV activity of efavirenz at low micromolar concentration (EC_50_ = 1.33 µM). To determine potential broad anti-alphavirus activity of efavirenz, its inhibitory activity against Sindbis virus (SINV) was detected. Interestingly, efavirenz also inhibited the replication of SINV at a low micromolar range (EC_50_ = 0.7 µM). Time of addition assay, direct transfection of virus replicon RNA and minus-sense specific RT-PCR elucidated that efavirenz hinders the viral replication at an early stage after the virus entry by inhibiting the viral RNA synthesis. Efavirenz showed a binding affinity with purified CHIKV capsid protein (CHIKV CP) and it is known that CHIKV CP plays a novel role in the early phase of viral protein synthesis, suggesting CP might be one of the potential protein targets in addition to viral or host proteins involved in viral genome replication. The present study underscores the repurposing of efavirenz for antiviral therapy against CHIKV to curb future viral outbreaks.

**Highlights:** Identification of *in vitro* anti-CHIKV activity of efavirenz.

Efavirenz disrupts the early phase of virus replication by interfering in the CHIKV minus-sense RNA synthesis.

Efavirenz and tomatidine also inhibit SINV replication indicating potential broad spectrum anti-alphavirus activity.

Efavirenz holds potential as therapeutic treatment for clinical infections of Chikungunya.

## 1. Introduction

Chikungunya virus (CHIKV), a re-emerging mosquito-transmitted alphavirus from the family *Togaviridae*, was first reported in Tanzania between 1952-1953 in the serum sample of a febrile patient (Robinson, 1955). CHIKV is an etiological agent of chikungunya fever characterized by polyarthralgia, headache, maculopapular rash, and asthenia (Bodenmann and Genton, 2006; Pialoux et al., 2007). CHIKV-infected patients experience chronic joint and musculoskeletal pain that might last months to years after post-acute infection, causing considerable suffering and economic loss (Suhrbier, 2019). From an epidemiological and medical perspective, CHIKV is the most prevalent alphavirus transmitted to humans by *Aedes albopictus* and *Aedes aegypti*. One hundred and fourteen countries over sub-tropical and tropical regions of Asia, Africa, Europe, and the Americas have reported autochthonous transmission of CHIKV (Puntasecca et al., 2021).

Earlier, the CHIKV outbreaks were limited to Sub-Saharan Africa (Powers and Logue, 2007); then, in 2004, the East/Central/South African (ECSA) strain re-emerged in Kenya (Chretien et al., 2007). The ECSA strain underwent rapid evolution and transmitted to new regions of islands in the Indian Ocean, India, and certain parts of Southeast Asia, resulting in an estimated 6 million cases (Powers and Logue, 2007; Staples et al., 2009). Within the clade ECSA, the Indian Ocean Lineage (IOL) displayed an alanine (Ala or A) to valine (Val or V) substitution mutation in the E1 glycoprotein at position 226. This adaptive mutation increased the transmission rate by 40-fold via the vector *Aedes albopictus* while not affecting viral fitness in the *Aedes aegypti* vector (Tsetsarkin et al., 2007; Volk et al., 2011). In addition to the mutation in CHIKV, the dense human population due to unplanned urban growth and modern rapid transportation has increased the risk of outbreaks (Ryan et al., 2018). Another major outbreak occurred in December 2013, when a strain from the Asian lineage emerged on Saint Martin Island in the Caribbean Sea (Cassadou et al., 2014). Almost every year, CHIKV outbreaks are reported around the globe (Bettis et al., 2022). Thus, to counter large-scale epidemics of CHIKV with high attack rates, there is an imminent need to develop effective therapeutics for outbreak containment.

The US Food and Drug Administration (FDA) recently approved Ixchiq (VLA1553), the first vaccine for chikungunya. However, vaccine-related serious side effects such as joint stiffness, muscle pain, fatigue, headache, fever and nausea were reported in less than 2% of the vaccinated population. The FDA needs a post-marketing study of Ixchiq to determine the risk of severe chikungunya-like adverse reactions (Ly, 2024). No specific licensed anti-CHIKV drug is available worldwide. As a result, the treatment primarily relies on symptom management using anti-inflammatory and analgesic medications for instance, paracetamol to alleviate the symptoms (Abdelnabi et al., 2015). However, targeting various non-structural proteins (nsPs) and structural proteins of CHIKV presents a promising avenue for finding effective drugs, as these viral proteins play a vital role in the replication cycle of the virus. Previous *in vitro* anti-CHIKV research has identified various compounds directed either toward nsPs, structural proteins of virus or host factors (Abdelnabi et al., 2015; Subudhi et al., 2018). To date chloroquine and the inosine monophosphate dehydrogenase inhibitor ribavirin have been tested in clinical trials (Lamballerie et al., 2008; Ravichandran and Manian, 2008). However, none of these compounds have been approved for treating human CHIKV infection, and the discovery of an effective drug treatment molecule has been unsuccessful (Kovacikova and van Hemert, 2020). Thus, developing effective anti-CHIKV drugs in clinical settings is imperative. Additionally, repurposing drugs already in clinical use for other diseases to target CHIKV viral proteins can offer a valuable strategy, potentially avoiding the need for the time-consuming and expensive drug trials of novel compounds (Pushpakom et al., 2018).

As all nsPs have some attributed and indispensable roles to play in the replication of viral genome (Lulla et al., 2006), hence nsPs are potential targets for developing effective antivirals (Mudgal et al., 2022, 2020; Pareek et al., 2022; Singh et al., 2018). The RNA genome of CHIKV comprises of two open reading frames (ORFs), of which the 5’ two-thirds of the RNA encodes for two overlapping polyproteins, P123 and P1234 (Khan et al., 2002; Solignat et al., 2009). The nsP2 protease processes these polyproteins into various intermediates and individual nsPs (Hardy and Strauss, 1989). These nsPs, with the help of host factors, contribute to architecting the replication complexes. First, the cleavage reaction forms the P123 and nsP4 (P123/nsP4) replication complex, which carries out the replication of minus-sense RNA by using genomic RNA as a template. Further, polyprotein P123 is cleaved at the 1/2 junction to form an nsP1/P23/nsP4 complex, which synthesizes both minus-sense and 49S genomic RNAs. Finally, fully processed nsPs are formed by cleavage at 2/3 junction of P23. Complete processed nsPs get involved in the synthesis of 26S subgenomic mRNA and genomic RNA (Shirako and Strauss, 1994; Vasiljeva et al., 2003).

In the present investigation, the anti-CHIKV activity of efavirenz has been identified, further potential broad anti-alphavirus activity was evaluated. Efavirenz is a non-nucleoside inhibitor of HIV and is currently in use for the treatment of acquired immunodeficiency syndrome (AIDS). Previously known anti-CHIKV compound tomatidine (Troost et al., 2020; Troost-Kind et al., 2021) was taken as a positive control, along with efavirenz; for the first time, this study reports the anti-SINV activity of tomatidine. Several approaches were applied to evaluate the possible mechanism of efavirenz anti-CHIKV activity. In total, this study indicate that efavirenz acts post-entry but in an early stage of CHIKV infection by affecting the functions of proteins involved in RNA replication and significantly inhibits the synthesis of minus-sense viral RNA. Furthermore, efavirenz interaction with CHIKV capsid protein (CHIKV CP) was observed. Thus, it is possible that along with affecting the functions of proteins involved in viral RNA replication, efavirenz might be inhibiting the functions of CP which are critical during the early phase of replication (Kiser et al., 2021).

## 2. Materials and Methods

A detailed description of the Materials and Methods is mentioned in the Supporting Information file.

## 3. Results

### 3.1. Identification of efavirenz as an inhibitor of CHIKV and SINV infection

Prior to evaluating the antiviral activity of compounds, the cytotoxicity of compounds was determined by performing 3-(4,5-dimethylthiazol-2-yl)-2,5-diphenyl tetrazolium bromide (MTT) assay. The half-maximal cytotoxic concentration (CC_50_) value for Vero cells were 53.62 and 17.19 µM for efavirenz and tomatidine, respectively. In the case of BHK-21 cells CC_50_ values were 40.91 and 23.84 µM for efavirenz and tomatidine, respectively (Table 1).

**Table 1:** The CC_50_, EC_50_, and selectivity index (SI = CC_50_ / EC_50_) values of efavirenz and tomatidine with respect to Vero and BHK-21 cell line.

| Compound | $CC_{50}$ (Vero cells) ($\mu\text{M}$) | $CC_{50}$ (BHK-21 cells) ($\mu\text{M}$) | CHIKV | | SINV | |
| --- | --- | --- | --- | --- | --- | --- |
| | | | $EC_{50}$ ($\mu\text{M}$) | SI ($\mu\text{M}$) | $EC_{50}$ ($\mu\text{M}$) | SI ($\mu\text{M}$) |
| Efavirenz | 53.62 | 40.91 | 1.33 | 40.35 | 0.72 | 74.47 |
| Tomatidine | 17.19 | 23.84 | 5.35 | 3.21 | 2.47 | 6.95 |

Anti-CHIKV and anti-SINV activity were determined by quantitative evaluation of progeny virus particles released in the supernatant of infected cells upon treatment of infected cells by 2-fold dilutions of compounds below cytotoxic concentration (Fig. 1). Efavirenz showed dose-dependent inhibition of CHIKV and SINV propagation at the 50 % effective concentration (EC_50_) of 1.33 µM and 0.7 µM, respectively (Fig. 1 A and C). The control compound tomatidine displayed potent inhibition of the CHIKV and SINV replication at the EC_50_ values of 5.35 µM and 2.47 µM, respectively (Fig. 1 B and D). The inhibition of CHIKV by tomatidine has already been reported (Troost et al., 2020; Troost-Kind et al., 2021); however, for the first time in this study tomatidine is investigated for its inhibitory activity against SINV replication. Therefore, the antiviral activities of efavirenz and tomatidine are not confined to CHIKV but also efficiently restrict the replication of other alphaviruses.

**Fig. 1:**
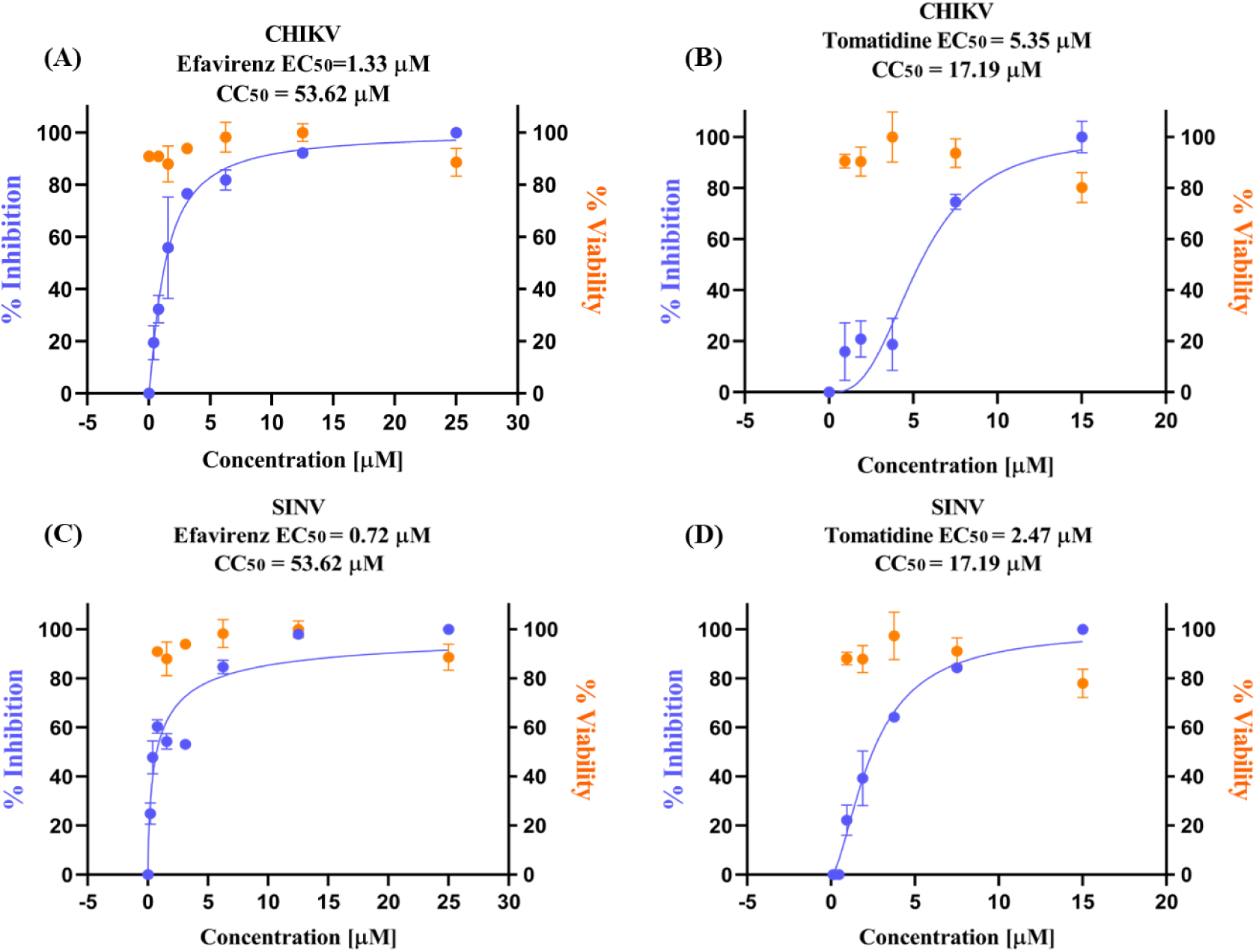
Efavirenz and tomatidine showed dose-dependent inhibition of released progeny virus particles of CHIKV (A and B) and SINV (C and D). The decrease in extracellular released CHIKV and SINV progeny virus particles titre 24 hours post-infection (hpi) in the presence of an increasing compound concentration was assessed by plaque assay. The percentage inhibition was calculated by considering 100% viral titre in the corresponding virus or vehicle control. The 50 % effective concentration (EC_50_) and half-maximal cytotoxic concentration (CC_50_) values were determined by GraphPad Prism 8 by using a non-linear fit model after data normalization. The values are the mean of two independent experiments and the error bar shows the standard deviation.

The intracellular replication inhibition of CHIKV and SINV was detected by Immunofluorescence assay (IFA) with the help of primary anti-alphavirus mouse monoclonal antibody and secondary FITC-conjugated antibodies. Reduction in intracellular fluorescence was observed for efavirenz and tomatidine treated cells (Fig. 2). Further, the reduction in the abundance of intracellular CHIKV RNA due to efavirenz and tomatidine treatment was detected by qRT-PCR. Significant fold change in the intracellular CHIKV RNA as compared to virus control was observed (Fig. 3). Thus, in addition to plaque assay based antiviral assay, IFA and qRT-PCR prove the excellent anti-CHIKV activity of efavirenz.

**Fig. 2:**
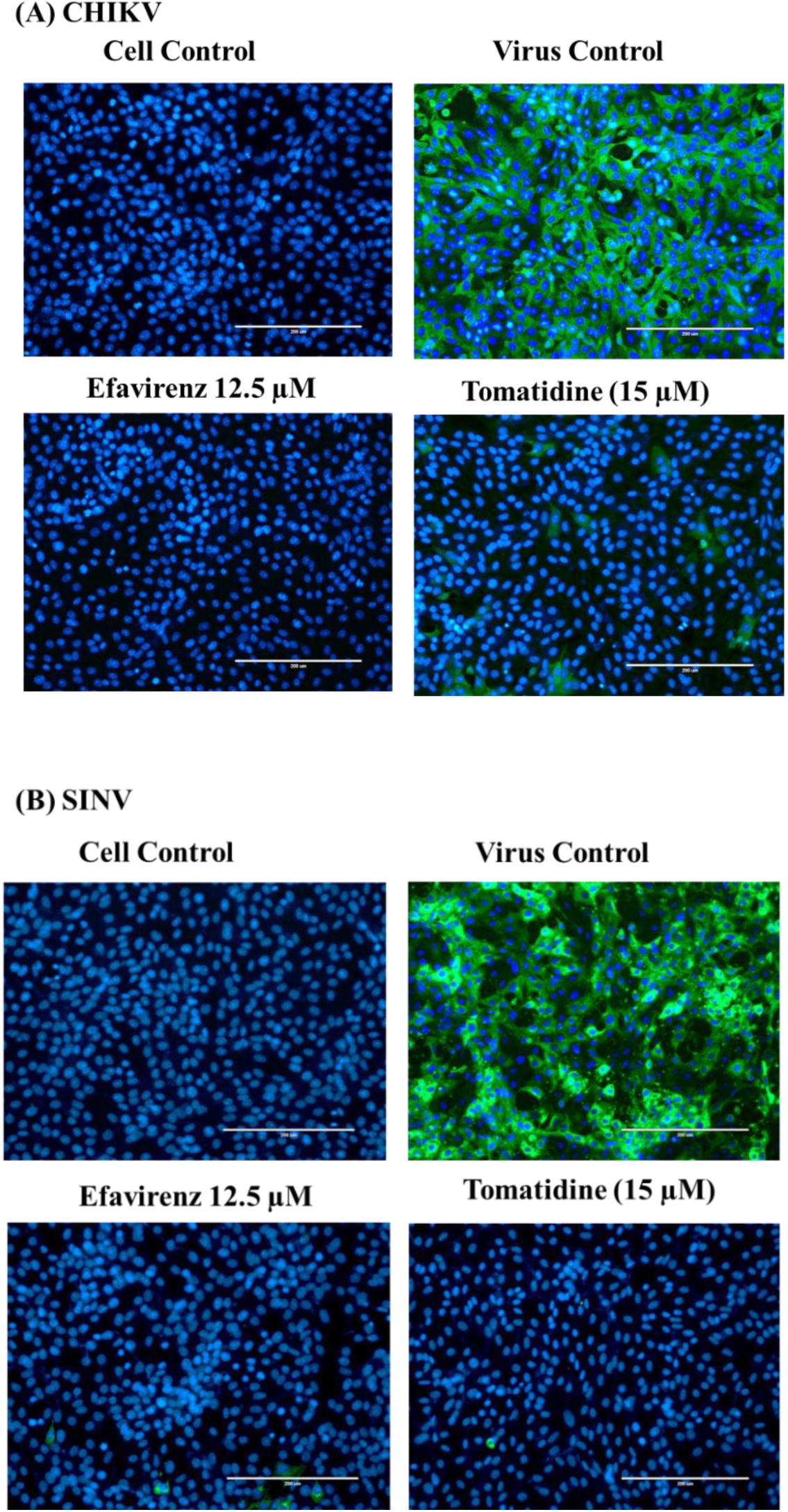
Detection of intracellular inhibition of A) CHIKV and B) SINV by efavirenz and tomatidine with the help of immunofluorescence microscopy. The cells were infected [Multiplicity of infection (MOI) = 1] and then treated with efavirenz (12.5 µM) and tomatidine (15 µM). After 30 hpi, the cells were fixed and the presence of replicated intracellular virus was detected by anti-alphavirus antibodies and fluorescein isothiocyanate (FITC)-conjugated secondary antibody (green). The 4’,6-diamidino-2-phenylindole (DAPI) was used to stain the cell nuclei (blue). The images were captured at 20 X magnification: scale bar, 200 µM.

**Fig. 3:**
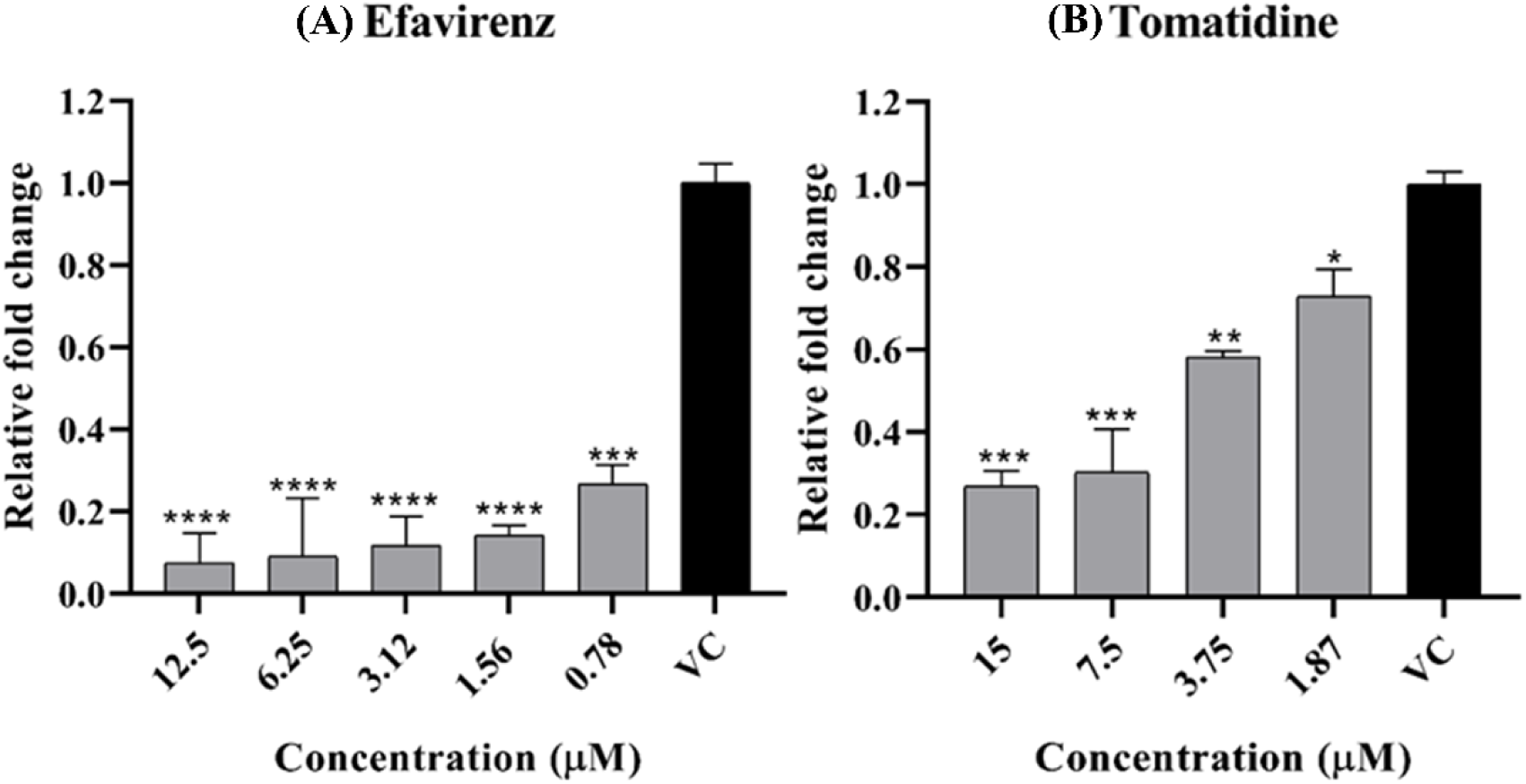
Relative fold change in the intracellular CHIKV RNA due to (A) efavirenz and (B) tomatidine treatment as compared to virus control (VC) was assessed by qRT-PCR. Total RNA was isolated 24 hpi and expression of E1 gene was assessed by E1 specific primers by keeping β-actin as an internal control. The data was analyzed by the one-way ANOVA test and Dunnett’s posttest; ****, P < 0.0001. ***, P = 0.0002. **, P = 0.0026, *, P = 0.0168. The error bar shows the mean values’ standard deviation (n = 2).

### 3.2. Evaluation of inhibition step of CHIKV life cycle by efavirenz

The time-of-addition (TOI) experiment was performed to find out the window when efavirenz exerts its anti-CHIKV activity. Vero cells were treated with efavirenz (6.25 µM) during varying time periods, such as 2 h before infection (Pre), simultaneous or at the time of infection (ATI) and post 0, 2, 4, 6, 8, 10, 12 h after infection (Fig. 4A). Efavirenz most prominently and significantly inhibited the CHIKV replication up to 6 h of post-infection (Fig. 4B). Pre and simultaneous or ATI treatment showed non-significant but some inhibition, indicating that some amount of efavirenz must be entered into cells and retained during the time of post-infection. Non-significant inhibition was observed when the compound was added 8 hpi, indicating a loss of efavirenz inhibitory activity. Thus, TOI experiment proved that efavirenz acts in an early phase of CHIKV replication after virus entry.

**Fig. 4:**
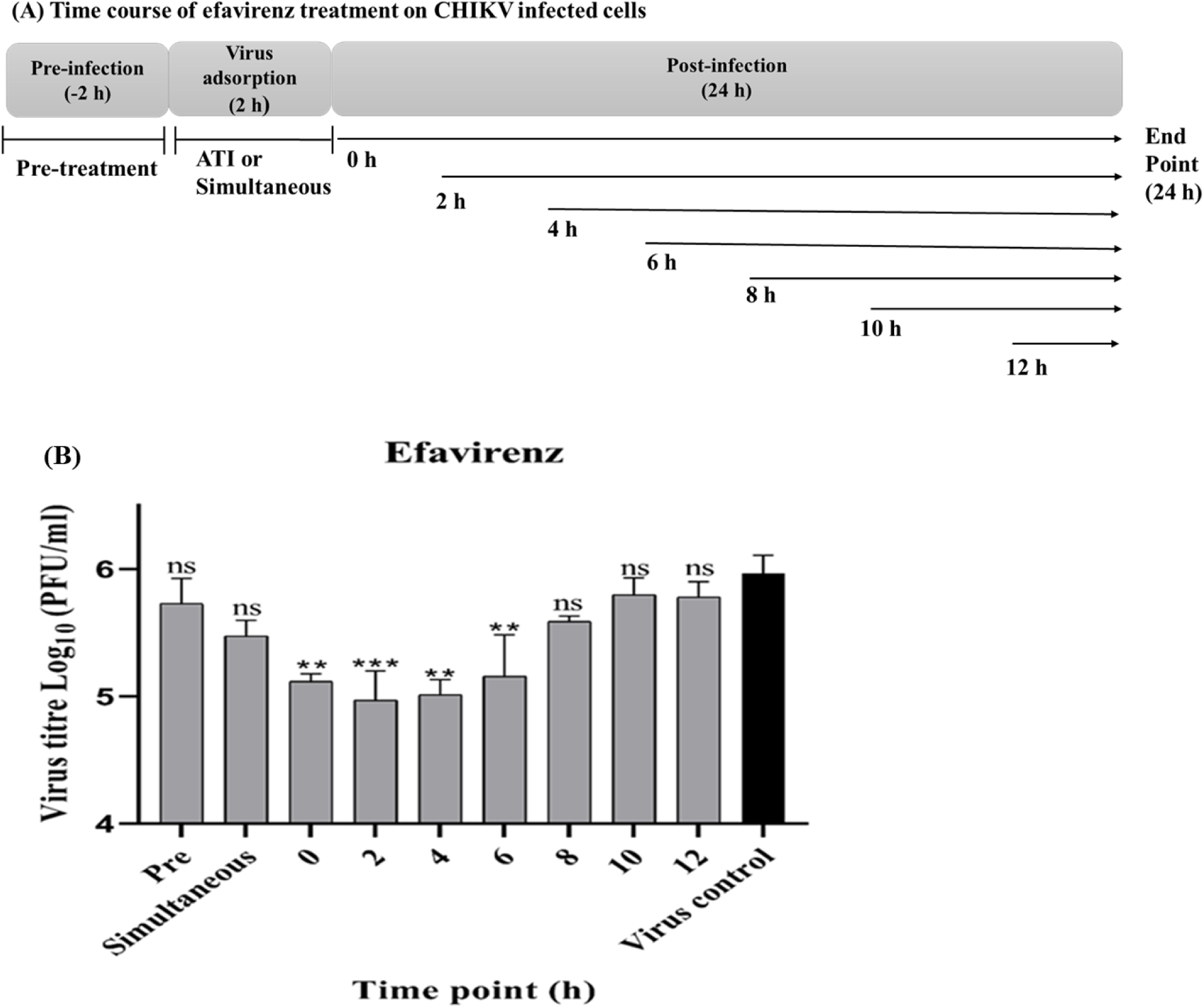
(A) Schematic representation of time course of efavirenz treatment on CHIKV infected cells. (B) The pronounced inhibitory effect of efavirenz is observed in the post-treatment of the compound up to 6 h of infection. Vero cells were infected with CHIKV (MOI = 0.1), and cells were treated before infection (–2 h), during infection and post-infection (0 to 12 h) at indicated time points. The data was analysed by the one-way ANOVA test and Dunnett’s test; ***, P = 0.0010. **, P = 0.0014 to 0.0046. ns, non-significant. The error bar shows the mean values’ standard deviation (n = 2).

### 3.2. Efavirenz and tomatidine modulates with the functions of non-structural proteins and downregulate the synthesis of viral minus-sense RNA

Remarkable anti-Sindbis virus replicon (SINV-REP) activity was observed when *in vitro* transcribed SINV-REP RNA was transfected into BHK-21 cells and further treated by efavirenz and tomatidine for 6 h (Fig. 5 B and C). Replicon assay involves direct transfection of viral RNA into cells, thus eliminating the virus particle binding with receptors and fusion steps with host cells. The tested compounds showed a significant inhibitory effect against SINV-REP, indicating suppression of viral replication machinery by compounds and further translation and expression of viral proteins. This result also ruled out the possible inhibitory effects of compounds on virus adsorption with the receptor or virus fusion step. However, the replicon assay did not entirely rule out the probability of efavirenz inhibitory activity against functions of structural proteins as more than one protein target of a compound is a possibility in the case of wild-type virus.

**Fig. 5:**
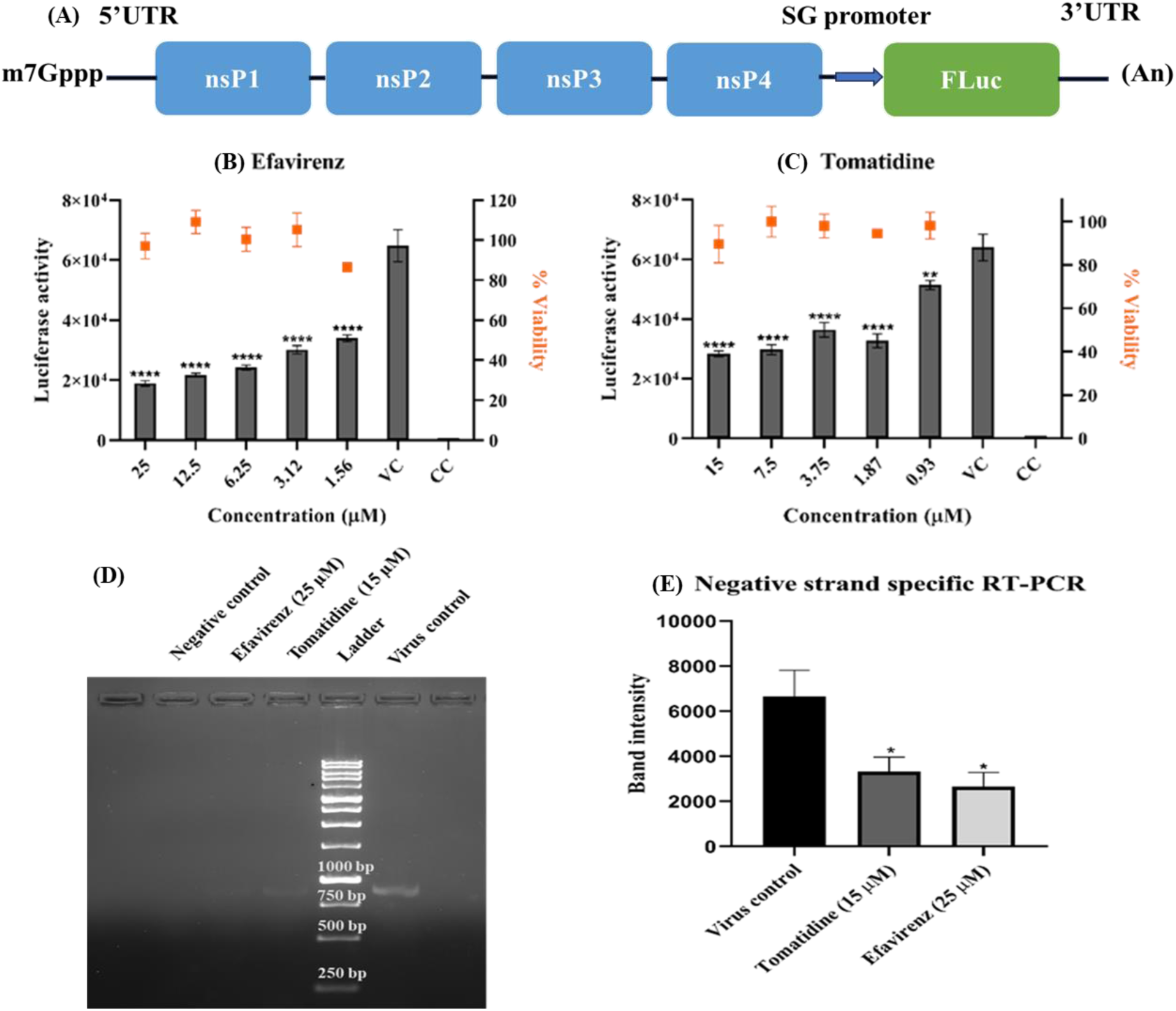
Efavirenz and tomatidine modulate the function of non-structural proteins and inhibit the synthesis of minus-sense viral RNA. (A) Schematic diagram of Sindbis virus replicon (SINV-REP). The SINV-REP is constructed by replacing the structural protein genes with Fluc downstream of SG promoter and IRES element at the 3’ end. Electroporated in vitro transcribed RNA of SINV-REP lacks the genes of structural protein thus, transfected capped RNA is only able to synthesize RNA replication complexes formed by non-structural proteins. (B and C) A significant decrease in luciferase activity was observed due to the treatment of increasing concentration of compounds efavirenz and tomatidine. (D and E) Efavirenz and tomatidine inhibit the synthesis of minus-sense viral RNA, the amplified products of minus-sense RT-PCR were run on 1.5 % agarose gel and visualized with the help of image lab software. The decrease in band intensity as compared to virus control was evaluated with the help of image lab software 6.1. The error bar represents the standard deviation from the duplicate experiment. The data was analyzed by the one-way ANOVA test and Dunnett’s test; ****, P < 0.0001. **, P = 0.0043. *, P = 0.0301 to 0.0488. The error bar shows the mean values’ standard deviation (n = 2).

The nsPs carry out synthesis of the viral RNA, thus effect of the compounds on the synthesis of viral minus sense RNA was evaluated by minus-sense specific RT-PCR. A significant decrease in the amplified band intensity was observed after treatment of CHIKV infected cells with efavirenz (25 µM) and tomatidine (15 µM) as compared to virus or vehicle control (Fig. 5 D and E). The amplified band are approximately in proportionate to the minus-sense viral RNA synthesised in the virus infected cells at the end of assay as the viral minus-sense specific E1 primer was used to synthesise cDNA and region of E1 gene was further amplified from cDNA for visualisation on the agarose gel. A marked decrease in band intensity indicates inhibition of the synthesis of minus-sense CHIKV RNA with the treatment of efavirenz and tomatidine. Thus, this experiment concludes that the compounds modulate the functions of nsPs or host factors which are involved in the synthesis of minus-sense viral RNA.

### 3.3. Biophysical characterization of efavirenz interaction with the non-structural proteins and structural protein

The previous experiments suggest that efavirenz is acting on nsPs or host factors to downregulate the minus-sense viral RNA synthesis. To investigate effect of efavirenz on nsPs as well as on CP, an attempt was made to identify the interaction of efavirenz with soluble, stable forms of CHIKV nsP1 (residue 1-509), CHIKV nsP2 (residue 471-791), and SINV nsP4 (N-terminal 97 residue truncated) and CHIKV CP (residue 106-261). These non-structural protein and CHIKV CP were expressed in the bacterial expression system and recombinant proteins were purified by Immobilized metal ion affinity chromatography (IMAC) (*Supplementary* Fig. 1) (Kaur et al., 2018; Pareek et al., 2022; Sharma et al., 2016; Singh et al., 2018). Isothermal titration calorimetry (ITC) and Surface plasmon resonance (SPR) are standard biophysical techniques for investigating the label-free molecular interactions of drug molecules with the target proteins. We investigated the affinity of efavirenz towards CHIKV nsP1 by ITC and towards CHIKV nsP2, SINV nsP4 and CHIKV CP by SPR in the range of 1 to 200 µM concentration. The efavirenz did not show detectable interaction with CHIKV nsP1, CHIKV nsP2, and SINV nsP4 (*Supplementary* Fig. 2). Efavirenz binding with CHIKV CP was observed with the K_D_ value of 6.22 µM (Fig. 6). Recent study of Kiser et al. has shown that CHIKV CP plays a novel role in regulation of viral protein synthesis processes early in the infection (Kiser et al., 2021). Thus, it is possible that inhibition of CHIKV and SINV, in addition to effect on RNA synthesis could be due to efavirenz interaction with CP. Tomatidine precipitated at concentrations higher than 15 µM; thus, binding experiments with the mentioned proteins could not be performed, limiting the ability to evaluate the interaction of tomatidine towards nsPs and CHIKV CP.

**Fig. 6:**
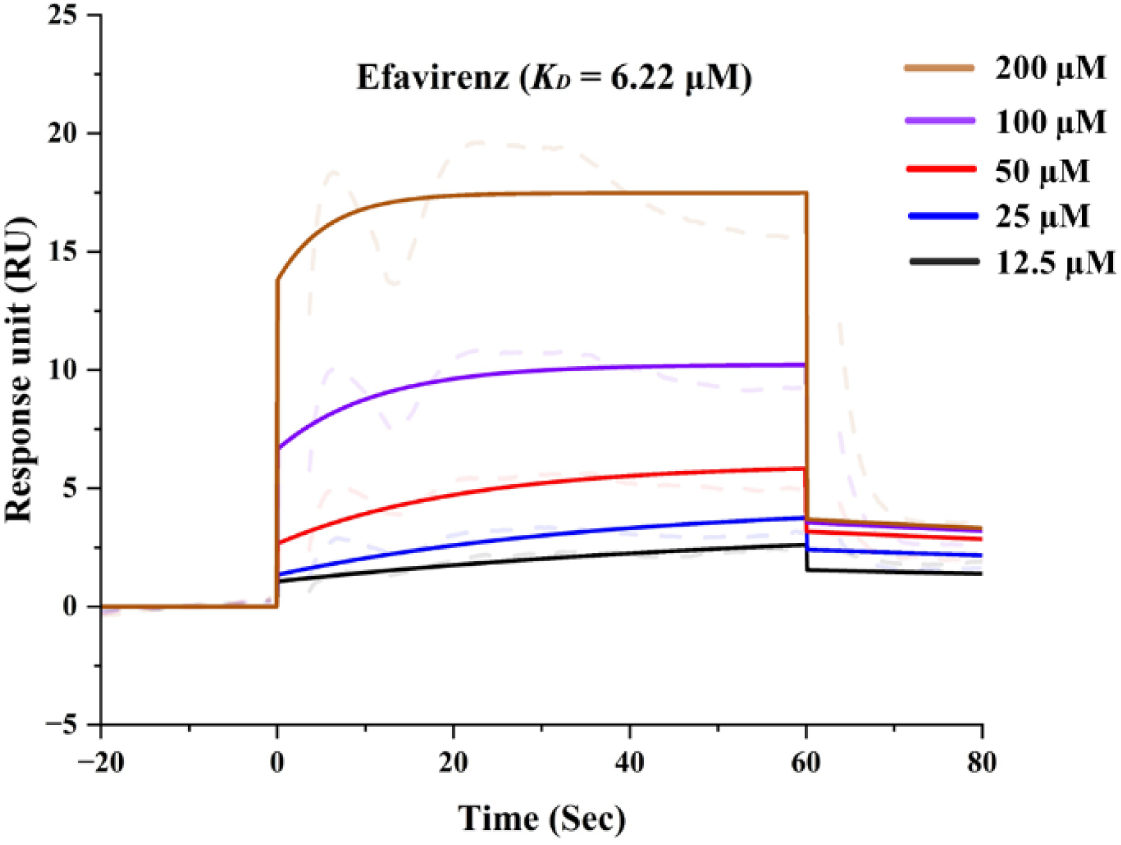
SPR sensorgram elucidating binding kinetics analysis of efavirenz onto the immobilized CP. The different colour-coded lines represent the varying efavirenz concentration (12.5 – 200 μM) injected onto the sensor surface of immobilized CP to detect the kinetic affinity. The positive binding with increasing concentration of efavirenz was observed with K_D_ Value of 6.22 µM.

## 4. Discussion

CHIKV is a recurring alphavirus with a global presence in tropical and sub-tropical regions that lacks effective licensed drugs. In our current study, the anti-alphavirus activity of efavirenz has been identified.

Efavirenz is a well-known inhibitor of HIV reverse transcriptase; it is the first-generation non-nucleoside reverse transcriptase inhibitor (NNRTI) used to treat acquired immunodeficiency syndrome (AIDS). Efavirenz is extensively used in the antiretroviral regimen to treat HIV-infected patients in developing countries because of its efficacy, affordability, low cost, and convenience (Nachega et al., 2008). The oral bioavailability of efavirenz is good and it has a long half-life. When a single oral dose of 100 to 1600 mg is administered to healthy volunteers, the peak efavirenz plasma concentrations of 1.6 – 9.1 µM reach after 5 hours of dose. On average, it takes 3 – 5 hours to reach peak plasma concentrations, and steady-state plasma concentrations can be reached in 6 – 10 days (Vrouenraets et al., 2007). Efavirenz has been reported to have antiviral activity against Zika and West Nile Virus (Sariyer et al., 2019; Stefanik et al., 2020). Previously, Varghase et al. had made an attempt to evaluate the anti-CHIKV activity of efavirenz by CHIKV replicons, but no significant inhibition was observed. However, Varghese et al. did not proceed to test anti-CHIKV activity on the wild type virus (Varghese et al., 2016).

In this study, the effective anti-CHIKV activity of efavirenz using an isolated clinical strain (Singh et al., 2018) has been demonstrated. The potential broad-spectrum anti-alphavirus activity of efavirenz has been proposed by demonstrating and detecting significant inhibition against SINV, another member of the alphavirus genus (Fig. 1). Robust inhibition in released infectious progeny virus particles was observed by plaque assay (Fig. 1), and a drastic reduction in intracellular viral replication was observed through IFA upon compound treatment compared to the virus control (Fig. 2 A and B). In the case of CHIKV, the abundance of intracellular RNA was assessed by qRT-PCR, and a significant reduction in intracellular CHIKV RNA copy number as compared to virus control (VC) was revealed upon compound treatment (Fig. 3). The efavirenz exhibited highly effective inhibition of CHIKV replication, with a low micromolar EC_50_ value of 1.33 µM (Fig. 1A) and also abrogated SINV replication (EC_50_= 0.7 µM) (Fig. 1C). A time of compound addition experiment was performed to understand better the window in which the CHIKV replication cycle is inhibited by efavirenz. The most potent inhibition of CHIKV was observed in the post-infection treatment up to 6 h of infection (Fig. 4B). Thus, the time of addition assay gave insights into the inhibitory mechanism of efavirenz against CHIKV, indicating that efavirenz is inhibiting an early step but post-binding and entry step of the virus replication cycle. During the early phase of infection, within the initial 6 h after entry into host cells, CHIKV RNA replication occurs (Albulescu et al., 2014). A time-based assay indicated that the efavirenz might be acting on the synthesis of viral RNA by either affecting the nsPs or host factors, which are vital for alphavirus RNA replication.

To evaluate the effect of efavirenz and the control drug tomatidine on the functions of alphavirus nsPs, we further employed the SINV-REP assay. SINV-REP lacks the structural protein genes and only has nsP genes; thus, it can be used to evaluate the effect of compounds on the functions of nsPs (Fig. 5A). A significant concentration-dependent decrease in luciferase activity with increasing concentration of compounds was observed, suggesting that the compounds exert the inhibition by modulating the functions of SINV nsPs (Fig. 5B and C). Strong inhibition of transfected RNA replication by compounds also excludes the inhibitory role of compounds during virus attachment to cells and entry, as these steps are bypassed. Along with previously performed antiviral assays in Vero cells, SINV-REP inhibition was observed in BHK-21 cells, thus demonstrating that the antiviral effect is not cell type-specific, therefore further strengthening the potential of efavirenz as drug compound to treat CHIKV infection. These results encouraged us to investigate the effect of compounds on the synthesis of minus-sense viral RNA which is exhibited by replication complex of nsPs. To gain further insight into the effect of compound treatment on the synthesis of alphavirus minus-sense RNA, the amount of minus-sense CHIKV RNA produced in the presence of compounds was compared with the virus or vehicle control with the help of minus-sense specific RT-PCR. A significant difference in the band intensity was observed upon compound treatment of CHIKV-infected cells (Fig. 5D and E). Minus sense-specific RT-PCR further narrowed down the possibilities of the inhibitory mechanism of both compounds by indicating that the compounds are affecting the synthesis of viral minus-sense RNA. The alphavirus RNA synthesis inhibition can be due to the direct effect of compounds on nsPs or indirectly by affecting host factors involved in viral RNA replication. Out of four nsPs, the N-7-methyltransferase activity of CHIKV nsP1 (residues 1 to 509) (Mudgal et al., 2020), the protease activity of nsP2 (residue 471-791) (Singh et al., 2018) and TATase activity of active core catalytic domain of the SINV nsP4 (N-terminal 97 residue truncated) (Pareek et al., 2022; Tomar et al., 2006) has been reported previously from our and other research groups. Among the four nsPs, the CHIKV nsP1, CHIKV nsP2, and SINV nsP4 expressed and purified from bacterial expression are soluble, stable and enzymatically active in aqueous buffer, thus can be utilized to check interaction with efavirenz with the help of biophysical techniques such as ITC and SPR. To evaluate the possible effect or interaction of efavirenz with CHIKV nsP1, ITC was used, whereas to detect interaction with CHIKV nsP2 and SINV nsP4, we employed the SPR technique within the efavirenz concentration range of 1 to 100 µM. Efavirenz has not shown interacted with CHIKV nsP1, CHIKV nsP2, or SINV nsP4. Therefore, the lack of affinity towards these proteins excludes the inhibition of nsP1 N-7-methyltransferase activity, nsp2 protease and nsP4 polymerase activity by efavirenz. Here, we have not performed the studies on interaction of efavirenz against full length of nsP1, nsP2 and nsP4 proteins as these could not be expressed in the bacterial expression system. Thus, efavirenz might be interacting with nsPs or host factors involved in viral RNA replication to exert its anti-CHIKV or anti-SINV activity. Hence, the exact protein target or targets by which efavirenz inhibits alphavirus RNA replication remains to be elucidated. To further investigate the inhibition mechanism of efavirenz in the early virus replication phase, the molecular binding study of the efavirenz with purified CHIKV CP was performed as CP plays a novel role in the early virus protein synthesis (Kiser et al., 2021). SPR study clearly demonstrated the binding of efavirenz with CHIKV CP with K_D_ values 6.22 µM (Fig. 6), and this CP-efavirenz interaction suggests that efavirenz seems to be inhibiting the role of CP in the early viral protein translation processes in addition to inhibition in viral RNA replication. Tomatidine was also selected for further ITC, SPR binding assay evaluation to determine its binding to the nsPs. However, due to low solubility above 15 µM concentration, the affinity of tomatidine towards the nsPs could not be evaluated with the help of ITC or SPR. Troost et al. reported EC_50_ values of tomatidine against different strains of CHIKV in the range of 1.3 to 3.8 µM. Compared to Troost et al. in our experimental settings, the obtained EC_50_ value of tomatidine was 5.35 µM. Further, Troost et al. have reported that tomatidine acts during the early phase of infection post-entry and controls the expression of viral proteins (Troost et al., 2020; Troost-Kind et al., 2021). Similarly, with different experimental approach, we also proved that tomatidine acts during early phase of viral infection by inhibiting the synthesis of minus-sense viral RNA after entry into the cells with the help of SINV-REP and minus sense-specific RT-PCR assay. In addition to this, anti-SINV activity of tomatidine was unknown. Here, we demonstrated the potential broad-spectrum anti-alphavirus activity of tomatidine by detecting inhibition of SINV propagation (EC_50_= 2.47 µM). It is noteworthy that tomatidine’s inhibitory effect extends beyond CHIKV, as it has also demonstrated activity against Zika and Dengue viruses (Diosa-Toro et al., 2019).

In summary, the study evaluated the anti-CHIKV activity of efavirenz and focused on elucidating the inhibition step in the virus replication cycle. Cell culture-based anti-viral assay, IFA, and qRT-PCR have demonstrated CHIKV replication abrogation by efavirenz. Potent broad-spectrum anti-alphavirus activity of both the compounds efavirenz and tomatidine was shown by significant inhibition of SINV replication. In addition to these findings, experimentally it was shown that both compounds control the synthesis of minus-sense viral RNA to exert antiviral activity by modulating the early replication processes. Further, the affinity of efavirenz against truncated non-structural and structural proteins was investigated. Binding was observed with the purified CHIKV CP thus, it is thus possible that one of the targets of efavirenz might be CP as it also plays an important role in the early viral protein translation of alphaviruses. Based on good pharmacokinetics properties, previous successful applications of efavirenz to treat HIV infection and demonstration of robust anti-alphavirus activity of efavirenz in the present study indicate excellent potential to repurpose efavirenz against CHIKV.

## Acknowledgement

This research was funded by ICMR (ISRM/12(46)/2020.) S.N. thank the Ministry of Human Resource Development (MHRD) and R.R thank the University Grants Commission (UGC), Government of India, for research fellowship. The authors also thank the Department of Biosciences and Bioengineering (BSBE), Government of India, for supporting Bioinformatics Centre at IIT Roorkee (reference number BT/PR40141/BTIS/137/16/2021). The authors acknowledge and thank the Macromolecular Crystallographic Unit (MCU), and Ashok Soota molecular medicine facility, Indian Institute of Technology, Roorkee.

## Author Contributions

**Shailly Tomar:** Conceptualization; Methodology; Project administration; Resources; Supervision; Validation; and Writing – original draft. **Sanketkumar Nehul:** Investigation; Methodology; Validation; Writing – original draft. **Ruchi Rani:** Investigation; Methodology; **Pravindra Kumar:** Project administration; Resources; Supervision; Writing – review & editing. **Prasan Kumar Panda:** Conceptualization; Writing – review & editing.

