## Supplementary Data for "Repurposing efavirenz, the HIV antiretroviral drug for Chikungunya virus infection"

### 1. Materials and methods

#### 1.1. Virus, cells, and compounds

The CHIKV patient sample isolate (Accession no. KY057363.1) (Singh et al., 2018) and full-length cDNA clone (pTOTO64) derived SINV were used in this study. Vero cell line (NCCS, Pune, India) was utilised for the propagation of viruses and evaluation of the antiviral activity of compounds, whereas the BHK-21 cell line (NCCS, Pune, India) was used for *in vitro* SINV replicon (SINV-REP) RNA transfection experiment. Both the cell lines were cultured in Dulbecco's modified eagle's medium (DMEM) (HiMedia, India) augmented with penicillin (100 U/mL), streptomycin (100 µg/mL) (HiMedia, India) and 10% heat-inactivated Fetal bovine serum (FBS) (Gibco, Brazil) at 37°C in the presence of 5% $CO_2$.

Efavirenz (E0997) was purchased from TCI Chemicals, India and tomatidine (T2909) was purchased from Sigma, India. The stock solutions of efavirenz and tomatidine were prepared in dimethyl sulfoxide (DMSO) (HiMedia, India) and stored at -20°C. Working dilutions of the compounds for treating the cells were prepared by diluting the stock solutions in DMEM supplemented with 2% FBS for cell culture-based assays and in 1X phosphate buffer saline (PBS) or 1X HEPES buffer saline (HBSN; 10mM HEPES, 150 mM NaCl, pH 7.4) for Isothermal titration calorimetry (ITC) or Surface plasmon resonance (SPR) based study.

#### 1.2. Cell viability assay

The viability of the Vero and BHK-21 cells in the presence of different compound concentrations was evaluated using 3-(4,5-dimethylthiazol-2-yl)-2,5-diphenyl tetrazolium bromide (MTT) (HiMedia, India) assay, as described previously (Fatma et al., 2020; Singh et al., 2018). Briefly, a 96-well plate containing Vero or BHK-21 monolayer was incubated with 2-fold dilutions of compounds in 2% FBS containing DMEM for 24 h at 37°C and 5% $CO_2$. After 24 h of compound treatment, 20 µL of MTT (5 mg/mL) was added per well. The cells were further incubated at 37°C in 5% $CO_2$ for 4 h. Upon incubation, formazan crystals were dissolved in DMSO and the absorbance was recorded at 570 nm using the plate reader (Multiskan Sky, Thermo Fisher Scientific). The average absorbance of cells treated with solvent was the control. The cell viability of the treated cells was compared to that of the control. The half-maximal cytotoxic concentration ($CC_{50}$) values were calculated using non-linear regression curve fit analysis.

#### 1.3. *In vitro* antiviral assay

To evaluate the anti-CHIKV and anti-SINV activity of compounds, a 24-well plate was seeded with Vero cells and incubated at 37°C / 5% $CO_2$ to form the monolayer. On the next day, the monolayer was infected by CHIKV or SINV with a 0.1 multiplicity of infection (MOI) while gentle shaking every 15 min for 1.5 h followed by washing with 1X PBS, and post-treatment with the 2-fold dilution of compounds in DMEM containing 2 % FBS was performed for 24 h. Virus control or vehicle control was incubated with 0.2 % DMSO in DMEM containing 2 % FBS to rule out the effect of solvent on viral replication. Upon incubation, 24 hours post-infection (hpi), the media was harvested and subjected to a plaque assay to quantify the number of progeny virus particles released in the supernatant. The 50% effective concentration ($EC_{50}$) value was calculated based on plaques forming units (PFU)/mL values obtained from the plaque assay by non-linear regression curve fit analysis. The percentage inhibition was calculated by considering 100% viral titre in corresponding vehicle-treated cells.

### 1.4. Immunofluorescence assay

A confluent monolayer of Vero cells in a 24-well plate was infected by 1 MOI of CHIKV or SINV for 1.5 h; subsequently, the monolayer was washed, and cells were incubated with efavirenz (15 µM) and tomatidine (12.5 µM) for 30 h at 37°C / 5% $CO_2$. After incubation, cells were fixed with a 1:1 ratio of methanol: acetone for 30 min followed by 1X PBS wash; subsequently, fixed cells were permeabilized by 0.1% Triton X-100 (Qualigens, India). Wash with 1X PBS was performed, and the anti-alphavirus mouse monoclonal antibody (Santa Cruz Biotechnology, USA) was incubated with cells for 90 min. Wash was given to remove excess amounts of primary antibody, and secondary fluorescein isothiocyanate (FITC)-conjugated anti-mouse secondary antibody (Sigma, USA) was added and incubated for 60 min. Finally, cells were counterstained using 4',6-diamidino-2-phenylindole (DAPI) (HiMedia, India) for 10 min. Infected cells treated with 0.2 % DMSO were considered virus or vehicle control and non-infected cells also treated with 0.2 % DMSO were taken as cell control. At the end of the assay, cells were observed under the EVOS FL imaging system (Thermo Fisher Scientific) using both DAPI and GFP channels. EVOS FL software was used to process the acquired images.

### 1.5. Intracellular CHIKV RNA assessment by quantitative reverse transcription polymerase chain reaction (qRT-PCR)

Total RNA was isolated from the cells after performing an *in vitro* antiviral assay by using Takara RNAiso Plus (Takara Bio, Japan) as per the manufacturer’s instructions.

Isolated RNA was quantified with the help of nanodrop, and 1μg RNA was used to synthesize cDNA by the PrimeScript first-strand cDNA synthesis kit (Takara Bio, Japan). E1 gene-specific primers were used as previously described and β-actin was kept as an internal control (Singh et al., 2018), using the KAPA SYBR fast universal qPCR kit on the QuantStudioTM 5 system (Applied Biosystems, USA). The qRT-PCR reactions were performed in duplicate, and the ΔΔCt method was used by calculating the relative quantification (RQ) value ($RQ = 2^{-\Delta\Delta Ct}$). The amplification in virus control was considered as 1 and the reduction in viral RNA expression in compound-treated cells was calculated as fold change by comparing it with the virus control.

### 1.6. Time of compound addition (TOA) experiment

Vero cells were treated with 6.25 μM concentration of efavirenz at pre-infection, simultaneous or at the time of infection (ATI) and post-infection with 0.1 MOI of CHIKV. For pre-treatment, efavirenz was added 2 h before the infection, and then cells were washed with PBS twice. For at the time of infection (ATI) condition, efavirenz was added to the virus inoculum and cells were incubated with efavirenz containing CHIKV for 2 h. In the case of post-treatment, cells were incubated with efavirenz at different time points of 0, 2, 4, 6, 8, 10, 12 h after infection. For all time point conditions, CHIKV infection was done for 2 h and the supernatant was harvested after 24 h of virus infection. The virus control was incubated with solvent (0.2 % DMSO) post-infection for 24 h.

### 1.7. Luciferase-based SINV replicon (SINV-REP) antiviral assay

SINV-REP assay was performed as described previously (Mudgal et al., 2022). *In vitro* RNA transcription was done from the clone pToto64 (Kind gift from Prof. Kuhn,

Perdue University). Capped RNA transcripts of SINV-REP were electroporated into BHK-21 cells in a 0.2 cm cuvette (25 μF, 1.5 kV, and 200 Ω) with the help of Gene Pulser Xcell (Bio-Rad). Then electroporated BHK-21 cells were seeded in the 24-well plate and allowed to attach for 4 h. Post-incubation, compounds or 0.2% DMSO (virus or vehicle control) were added to the transfected cells, and cell lysate was collected 6 h after the addition of compounds. The luciferase activity of cell lysates was measured using a firefly luciferase assay kit (Promega, USA) according to the manufacturer's instructions.

#### 1.8. Detection of compound treatment effect on CHIKV minus-sense RNA synthesis

To detect the levels of CHIKV minus-sense RNA in infected cells after compound treatment, minus sense-specific reverse transcriptase PCR (RT-PCR) was performed as described previously (Mayuri et al., 2008; Pareek et al., 2022). Confluent monolayer of Vero cells was treated 2 h prior to infection with 25 μM of efavirenz and 15 μM of Tomatidine in DMEM plus 2 % FBS. After 2 h of pre-treatment by compounds, compounds were removed and monolayer was washed with 1 X PBS, followed by CHIKV infection (10 MOI) for 1.5 h. Subsequently, cells were washed with 1X PBS and incubated with efavirenz (25 μM) and tomatidine (15 μM) for 6 h at 37°C, 5 % $CO_2$. At the end of 6 h, media was discarded and total cytoplasmic RNA was isolated by Takara RNAiso Plus (Takara Bio, Japan) and treated with DNase I, RNase-free (ThermoFisher, USA) as per the manufacturer's instructions. The quality and quantity of isolated RNA was detected by Nanodrop and further 1 μg of RNA was used to synthesis minus strand specific cDNA by using plus-sense primer (5'-GGAAATAACATCACTGTAACTGCCTATGCAAACG-3') and PrimeScript first-strand cDNA synthesis kit (Takara Bio, Japan). One-tenth diluted cDNA was further used for PCR reaction with the minus sense primer (5'-GCATAGCACCACGATTAGAATC-3') for 30-cycle amplification (Singh et al., 2018). The amplified products were run on 1.5 % Tris-Acetate-EDTA agarose gel containing ethidium bromide stain. The amplified band of 914 bp was visualized by the ChemiDoc MP Image system (Bio-Rad) using Image Lab Software 6.1 (Bio-Rad). The levels of minus-strand RNA synthesis was evaluated by analyzing the intensity of the bands of RT-PCR products on 1.5% agarose gel with the help of Image lab software 6.1.

### 1.9. Expression and purification of alphavirus proteins

#### 1.9.1. CHIKV nsP1

The CHIKV nsP1 (residues 1–509) was purified as described by Ramanjit Kaur et al. (Kaur et al., 2018). The secondary culture (2L) of Rosetta cells transformed with pET28c carrying nsP1 gene was induced with 0.4 mM isopropyl-b-1-thiogalactopyranoside (IPTG) at 18°C/180 RPM. At 16 h after induction, cells were harvested and lysed in buffer 50 mM HEPES, pH 7.3, 200 mM NaCl, 5% glycerol, and 20 mM Imidazole at 18 KPSI by French press (Constant Systems Ltd, England). The soluble protein present in the supernatant was purified by Immobilized metal-assisted chromatography (IMAC). The protein eluted in buffer 50 mM HEPES pH 7.3, 250 mM imidazole, 5% glycerol, 200 mM NaCl was analyzed by 12 % sodium dodecyl sulfate-polyacrylamide gel electrophoresis (SDS-PAGE). After dialysis against the 25 mM HEPES pH 7.3, 200 mM NaCl, the protein was concentrated with the help of an Amicon concentrator (cut off 30 kDa). Concentrated protein was further used for the ITC experiment to evaluate affinity towards efavirenz.

#### 1.9.2. CHIKV nsP2

The CHIKV nsP2 (residue 471-791) was purified as described previously (Singh et al., 2018). The pellet obtained from 1 L secondary recombinant bacterial culture induced by 0.4 mM IPTG (Himedia, India) was resuspended in 40 mL lysis buffer (50 mM Tris, pH 7.5, 500 mM NaCl, 10 mM imidazole and 5% glycerol). Resuspended cells were lysed with the help of the French press. Cell debris was removed by centrifugation, and the soluble protein present in the supernatant was purified using IMAC. The nsP2 was eluted in elution buffer (50 mM Tris buffer, pH 7.5, 250 mM NaCl, 300 mM imidazole and 5% glycerol) and elution fractions containing pure nsP2 were analyzed on 12% SDS-PAGE. Pooled elution fractions were dialyzed against 1X PBS at 4 °C overnight; further dialyzed protein was used for the SPR study.

#### 1.9.3. SINV nsP4

Previously published protocol for expression and purification of N-terminal 97 residue truncated nsP4 RdRp domain (SINV D97nsP4) was followed to obtain the pure SINV nsP4 protein (Pareek et al., 2022; Tomar et al., 2006). Briefly, pellet obtained from IPTG induced secondary 1 L recombinant bacterial culture was resuspended in lysis buffer (20 mM Tris-HCl, 40 mM imidazole, and 300 mM NaCl, pH 7.8) and subsequently, cells were lysed with the help of French press at 18 KPSI. The supernatant was loaded on the Ni-NTA beads for purification by IMAC. Further, to remove impurities, a wash with 1 M NaCl was given. Finally, the protein was eluted with 400 mM imidazole in buffer 20 mM Tris-HCl, pH 7.8, 300 mM NaCl. The elution fractions were analyzed by running the samples on SDS-PAGE.

Pure fractions of nsP4 were pooled and dialysed against 1X HBSN buffer saline. After overnight dialysis at 4°C, the pure protein was utilized by SPR assays to detect interaction with efavirenz.

#### 1.9.4. CHIKV capsid

The CHIKV capsid (CHIKV CP) was expressed and purified as previously described by Sharma et al. (Sharma et al., 2016). The recombinant construct pET28c containing the CHIKV CP gene and N-terminal 6X-his-tag was transformed into *Escherichia coli (Rosetta)*. When the OD600 was around 0.6, the culture was induced with 0.4 mM isopropyl-β-D-1-thiogalactopyranoside (IPTG) and further incubated for five h at 37 °C with continuous shaking at 180 rpm. Post incubation, the culture was harvested by centrifugation. For purification, the culture pellet was resuspended in 50 mL lysis buffer containing 50 mM Tris-HCl (pH 7.6), 100 mM NaCl, and 10 mM imidazole. Cells were lysed by a French press (Constant Systems Ltd, Daventry, England). After centrifugation, the soluble protein in the supernatant was purified by IMAC. The purified fractions containing CHIKV CP were pooled and dialyzed in 1X PBS at 4 °C for overnight and analyzed using 12% sodium dodecyl sulfate (SDS)-polyacrylamide gel electrophoresis (PAGE). Following dialysis, the protein was further utilized for the SPR.

### 2. Biophysical evaluation of efavirenz interaction towards non-structural proteins

The binding affinity of efavirenz against the purified CHIKV nsP1 was evaluated by MicroCal ITC-200 (GE Healthcare). Affinity towards CHIKV nsP2, SINV nsP4 and CHIKV CP was investigated using SPR by a Biacore T200 system (GE Healthcare).

### 2.1. Isothermal Titration Calorimetry (ITC)

ITC was performed to detect and calculate the possible binding thermodynamic parameters between CHIKV nsP1 and efavirenz on MicroCal ITC-200 (GE healthcare). The purified CHIKV nsP1 and efavirenz were diluted into buffer 25 mM HEPES pH 7.3, 200 mM NaCl. The efavirenz (100 µM) titration into 10 µM concentration of CHIKV nsP1 was performed at 25°C at set parameters described in Table 2. The obtained data was analyzed by MicroCal origin 7.0 software.

*Table 2: Different set parameters to perform the isothermal titration calorimetry experiments to characterize the binding of CHIKV nsP1 and efavirenz.*

| **Parameters** | |
|---|---|
| Total number of injections | 20 |
| Cell temperature | 25°C |
| Reference power | 8 |
| Initial delay | 120 s |
| Syringe concentration | 100 µM (Efavirenz) |
| Cell concentration | 10 µM (CHIKV nsP1) |
| Stirring speed | 750 rpm |
| Volume of 1st Injection | 0.5 µl |
| Duration of 1st injection | 1 s |
| Volume after 1st injection | 2 µl |
| Duration after 1st injection | 4 s |
| Injection spacing | 120 s |

| | |
|---|---|
| Filter period | 5 s |

### 2.2. Surface Plasmon Resonance (SPR)

The Ni-reagent kit (GE Healthcare) method was used to immobilize proteins over the Ni-NTA sensor chip (GE Healthcare). To prepare the Ni-NTA sensor chip, degassed and filtered 1X PBS or HBSN running buffer was used for pre-conditioning prior to immobilization of CHIKV nsP2 or SINV nsP4 or CHIKV CP. The purified his-tagged CHIKV nsP2 (10 µg/mL) or SINV nsP4 (10 µg/mL) or CHIKV CP (10 µg/mL) was immobilized on Ni-NTA sensor chips. The compounds were diluted in 1X PBS or HBSN buffer and subsequently flowed over the surface of immobilized protein at a 30 µl/min flow rate. The contact time for compound-protein interaction was set to 60 s, followed by a dissociation time of 60 s. The surface was regenerated after each dissociation phase using 350 mM ethylenediamine tetraacetic acid (EDTA) (HiMedia). The SPR data were acquired and subsequently analyzed using the Biacore evaluation software.

### 2.3 Statistical analyses

All statistical analysis were performed by GraphPad Prism 8. The 50 % effective concentrations ($EC_{50}$) and half-maximal cytotoxic concentrations ($CC_{50}$) were calculated by a non-linear fit model after data normalization. A one-way analysis of variance (ANOVA) test, a Dunnett's posttest, was implemented to determine the compounds' concentrations, resulting in a statistically significant difference compared to the Virus control or solvent control (0.2 % DMSO).

### 3. Results

#### 3.1. Purification of non-structural and structural proteins:

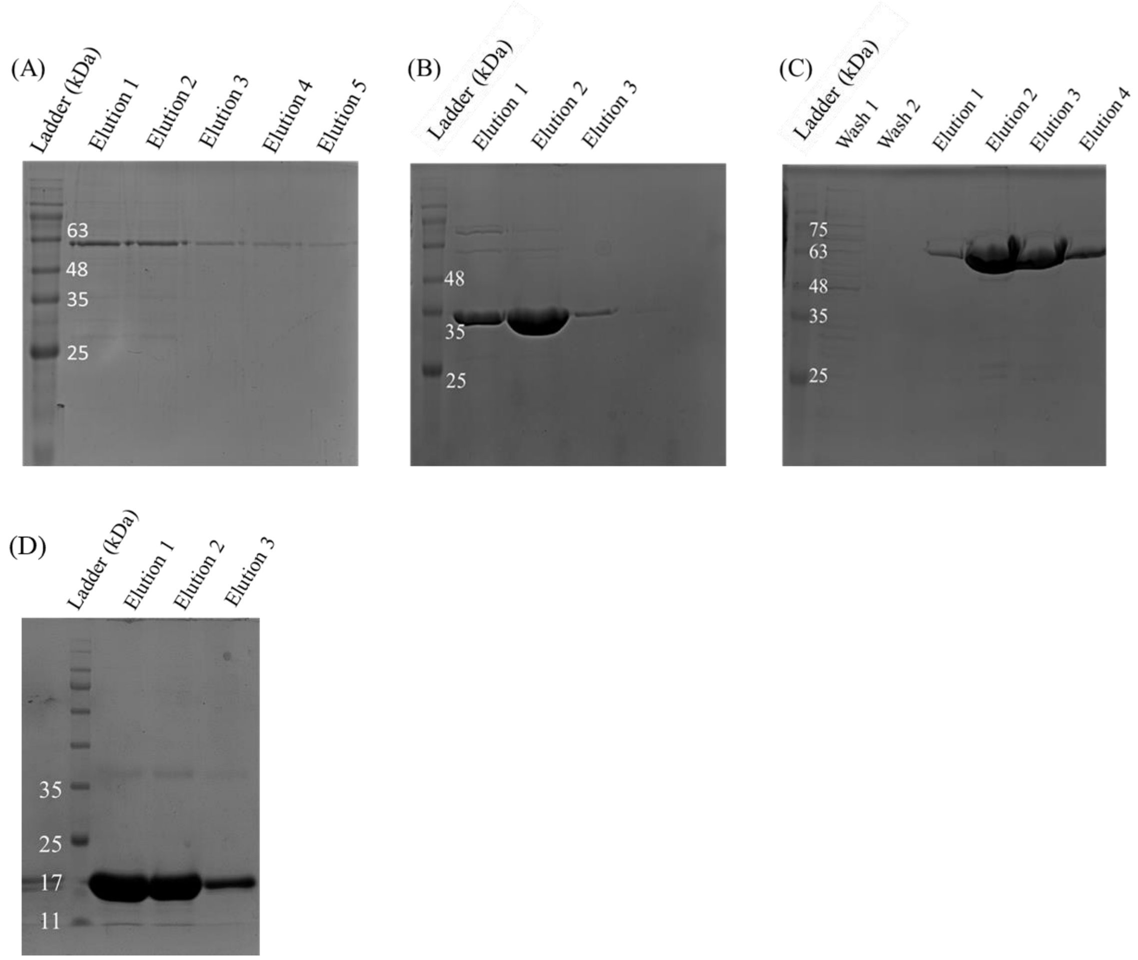


***Supplementary fig. 1***: *SDS-PAGE analysis of purified (A) CHIKV nsP1 (~ 56 kDa, residues 1–509), (B) CHIKV nsP2 (~ 35 kDa, residue 471-791), (C) SINV nsP4 (~ 63 kDA, N-terminal 97 residue truncated) and (D) CHIKV capsid (~ 18 kDa, residue 106-261) proteins. The proteins were expressed in the bacterial expression system and purified by IMAC, further elution fractions were pooled and proteins were dialyzed against buffer 1X PBS or 1X HBSN.*

#### 3.2. Evaluation of efavirenz interaction with non-structural proteins by biophysical techniques

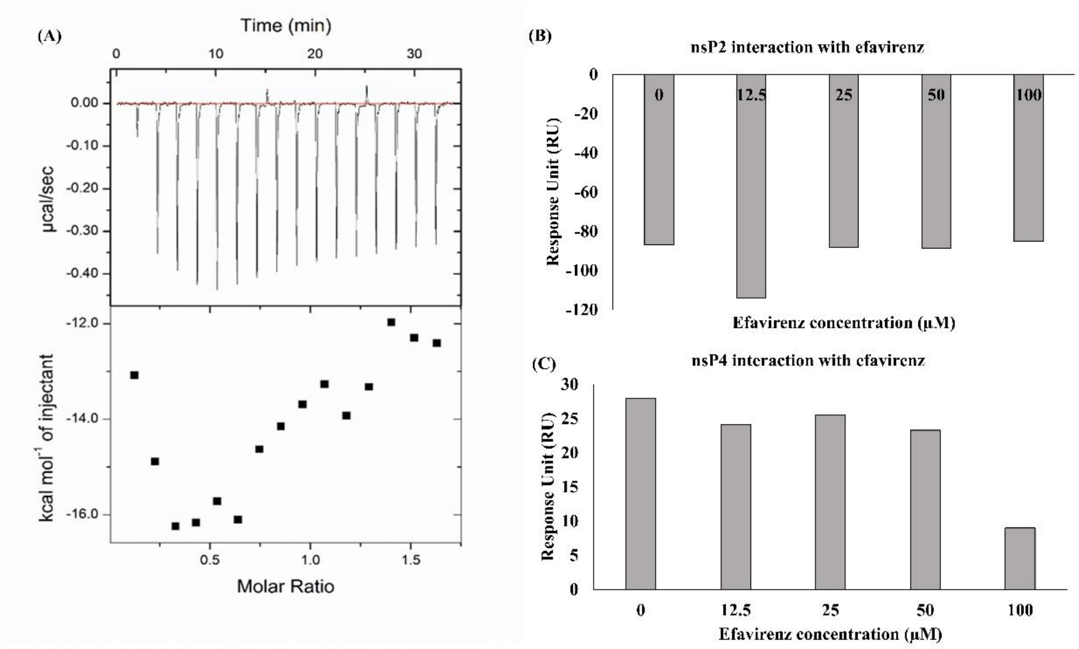


***Supplementary fig. 2:*** *Biophysical analysis of efavirenz interaction with non-structural proteins (A) Calorimetric titration of 15 injections each of 2 µl efavirenz (100 µM) into CHIKV nsP1 (10 µM) at 25°C. Negligible heat change without saturation shows lack of affinity between CHIKV nsP1 and efavirenz. (B) The purified CHIKV nsP2 and (C) SINV nsP4 proteins were immobilised on Ni-NTA chip and various concentrations of efavirenz (0 to 100 µM) were flown on immobilised chip to detect the interaction. A response unit (RU) increase in proportion to increased efavirenz concentration was not observed for CHIKV nsP2 and SINV nsP4, indicating an absence of interaction.*
